# IL-1RA Disrupts ATP Activation of P2RX7 in Human Monocyte-Derived Microglia-like Cells

**DOI:** 10.1101/2024.04.08.588607

**Authors:** Kelsey Heavener, Khushbu Kabra, Maedot Yidenk, Elizabeth Bradshaw

## Abstract

The immune system has a dynamic role in neurodegenerative diseases, and purinergic receptors allow immune cells to recognize neuronal signaling, cell injury, or stress. Purinergic Receptor 7 (P2RX7) can modulate inflammatory cascades and its expression is upregulated in Alzheimer’s disease (AD) brain tissue. P2RX7 expression is enriched in microglia, and elevated levels are found in microglia surrounding amyloid-beta plaques in the brain. While P2RX7 is thought to play a role in neurodegenerative diseases, how it modulates pathology and disease progression is not well understood. Here, we utilize a human monocyte-derived microglia-like cell (MDMi) model to interrogate P2RX7 activation and downstream consequences on microglia function. By using MDMi derived from human donors, we can examine how human donor variation impacts microglia function. We assessed P2RX7-driven *IL1β* and *IL18* production and amyloid-beta peptide 1-42 (Aβ1-42) uptake levels. Our results show that ATP-stimulation of MDMi triggers upregulation of *IL1β* and *IL18* expression. This upregulation of cytokine gene expression is blocked with the A740003 P2RX7 antagonist. We find that high extracellular ATP conditions also reduced MDMi capacity for Aβ1-42 uptake, and this loss of function is prevented through A740003 inhibition of P2RX7. In addition, pretreatment of MDMi with IL-1RA limited ATP-driven *IL1β* and *IL18* gene expression upregulation, indicating that ATP immunomodulation of P2RX7 is IL-1R dependent. Aβ1-42 uptake was higher with IL-1RA pretreatment compared to ATP treatment alone, suggesting P2RX7 regulates phagocytic engulfment through IL-1 signaling. Overall, our results demonstrate that P2RX7 is a key response protein for high extracellular ATP in human microglia-like cells, and its function can be modulated by IL-1 signaling. This work opens the door to future studies examining anti-IL-1 biologics to increase the clearance of amyloid-beta.

## Background

Alzheimer’s disease is a neurodegenerative condition characterized by continuous cognitive decline impacting a patient’s mood, memory, and ability to carry out daily tasks. This neurodegenerative disorder is the most common cause of dementia worldwide, and cases are expected to climb continuously as the global population ages.^1^ Currently, therapeutic options for slowing the progression of AD are limited, and no treatment options to prevent or reverse the diseased state exist.

AD pathology is defined by extracellular amyloid beta plaques and tau neurofibrillary tangles (NFTs) in the brain that are predicted to drive cognitive dysfunction.^2,3^ These aberrant proteins disrupt cell-to-cell communication and function, ultimately creating a neurotoxic environment.^4,5^ After decades of research centered around how amyloid beta and tau interact with neurons, there is now rising interest in studying the immune system’s contribution to AD. The primary drivers of the central nervous system (CNS) immune response are microglia. These resident immune cells of the CNS sample their environment and propagate differential immune responses to pathogens or injury including cytokine release, cell migration, and phagocytosis.^6^ Any pathogen presence or tissue injury will be detected by microglia, which become activated, drive inflammatory responses, and alert other cells nearby.

Microglia can detect the presence of amyloid beta and quickly migrate toward plaques.^7^ Microglia are found to congregate around amyloid beta plaques in both the human AD brain and mouse models of amyloidosis.^8,9^ As the professional phagocytes of the brain, microglia engulf amyloid-beta from their environment to maintain homeostasis in the brain.^10,11^ It has been postulated that these phagocytic functions are impaired in AD microglia due to proinflammatory signaling and/or aging and senescence.^12,13^ Removal of amyloid-beta accumulation in the brain has long been examined as a pathway for AD therapeutics and disease prevention.

One mechanism through which microglia survey their extracellular environment is through purinergic receptors. Purinergic receptors can bind extracellular ATP (eATP) outside the cell and become activated.^14,15^ The role of eATP in neurodegenerative disorders is an area of active investigation. Receptor activation through ATP triggers various downstream cellular functions including inflammation and apoptosis.^16,17^ In homeostatic, healthy conditions, eATP is found at low levels outside of the cell. However, when high ATP concentrations exist outside the cell, ATP functions as a damage-associated molecular pattern signal (DAMP). DAMPs are primarily endogenous molecules that, when released and detected outside the cell, signal imminent damage, and call the immune system to action. Apoptotic cells can release DAMPs, but they can also be transported into the extracellular space by vesicles.^18^ ATP can be released from distressed cells even prior to or without cell death, making it an important communication molecule for both tissue injury and homeostatic conditions.^19^ In response to eATP, microglia migrate to the site of injury or toxicity.^20^ It is likely these ATP-release avenues act in combination to create the elevated eATP conditions needed to activate P2RX7.

P2RX7, formerly classified as P2Z, is an ATP-gated ionotropic purinergic receptor that modulates the microglia phenotype in response to eATP.^21^ P2RX7 is highly expressed in immune cells, and expression is particularly enriched in microglia.^22–24^ This receptor is functionally distinct from its P2X_1-6_ family members in that it becomes activated by much higher concentrations of ATP.^25^ Upon chronic high eATP stimulation, P2RX7 forms a large membrane pore admitting molecules up to 900 Da in a non-selective manner.^26,27^ P2RX7 functions as an ion channel propagating Na^+^ and Ca^2+^ influx and K^+^ efflux. This ion movement modulates downstream signaling within the cell affecting cell migration and cytokine release.^28–30^ The Ca^2+^ influx mediated by P2RX7 also controls neurotransmitter release.^31^ These traits allow P2RX7 to modulate physiological intracellular processes in both homeostatic maintenance and proinflammatory function.

P2RX7 upregulates expression and production of cytokines including Interleukin-1 beta (IL-1β), Interleukin 6 (IL-6), and tumor necrosis factor α (TNF-α) in rats.^32^ Deletion of P2RX7 prevents IL-1β production and cytokine cascade in mouse macrophages.^33^ This cytokine release is partially mediated by activation of NLRP3 (NOD-,LRR-and pyrin domain-containing protein 3) inflammasome upon P2RX7 activation. Extracellular ATP binds to and activates P2RX7, triggering NLRP3 inflammasome activation.^34^ Experiments in human macrophages have demonstrated that ATP-mediated activation of P2RX7 results in NLRP3-inflammasome activation and IL-1β release.^35^ P2RX7 can directly colocalize with NLRP3 and trigger cytoplasmic ion changes that activate NLRP3.^36^ NLRP3 then recruits Caspase-1 which undergoes self-cleavage and activation. Activated Caspase-1 cleaves pro IL-1β and Interleukin 18 (IL-18) into their mature forms which are released from the cell.^37^ IL-18 expression is elevated in the brains of AD patients.^38^ Increased serum IL-1β levels have been observed in patients with mild cognitive impairment and AD.^39^ NLRP3 inflammasome is implicated in a range of neurodegenerative conditions including AD, Parkinson’s disease (PD), and traumatic brain injury.^40^ Hyperactivation of P2RX7 in the COVID-19-induced high eATP brain environment and subsequent NLRP3 inflammasome activation was proposed as a mediator of COVID-19 neuropathology.^41^

IL-1β is a powerful inflammatory cytokine produced by immune cells. It is a pivotal driver of inflammatory responses to injury and pathogens.^42^ The interleukin receptor antagonist protein (IL1RA) is a naturally occurring inhibitor of IL-1β initially discovered in serum and urine of children with chronic arthritis.^43^ By binding to the interleukin 1 receptor (IL-1R), IL1RA blocks IL-1α and IL-1β from binding to their target. This competitive inhibition prevents IL-1α and IL-1β from eliciting downstream inflammatory pathways.^44^

P2RX7 is implicated in a wide range of therapeutically challenging neurodegenerative disorders including Amyotrophic lateral sclerosis (ALS), Multiple Sclerosis (MS), PD, and AD. In both ALS and MS spinal cords, elevated P2RX7 levels were found in activated microglia.^45^ Chronic treatment of SOD1-G93A mice with the P2RX7 antagonist JNJ-47965567 improved motor performance and delayed disease onset.^46^ In MS, two gain of function SNPs in P2RX7 are associated with more severe disease.^47^ In contrast, a different rare P2RX7 variant that results in some loss of the pore function conferred a protective effect on risk for MS.^48^ In experimental autoimmune encephalomyelitis (EAE), which serves as a model of MS, P2RX7 deficiency suppressed disease development in mice.^49^ Blocking P2RX7 and its related ATP excitotoxicity in EAE also led to a reduction in demyelination and less damage to axons.^50^ P2RX7 is also implicated in PD pathways and related pathologies. In a Chinese population study, a P2RX7 SNP thought to influence receptor channel pore formation was associated with sporadic PD.^51^ Upregulation of P2RX7 has been found in the brains of human patients with tauopathies.^52^ Absence of P2RX7 in the THY-Tau22 mouse model of Tauopathy reduced the presence of inflammation and microglial activation, as well as led to improvements in synaptic plasticity and cognition.^52^

In AD, elevated P2RX7 levels are found at both the gene expression and protein level in mouse models and AD patients.^53,54^ Microglia derived from AD patients showed upregulation of P2RX7, particularly in association with Amyloid beta plaques.^54^ In sequencing experiments using 5XFAD transgenic mouse brains P2RX7 was upregulated in the disease-associated microglia (DAM) subtype.^55^ In an APP/PS1 mouse model, P2RX7 expression in microglia was significantly associated with amyloid beta-induced loss of postsynaptic density.^56^ In primary mouse microglia, where small interfering RNA (siRNA) was used to silence P2RX7, microglial phagocytosis of amyloid beta, was upregulated.^57^

A study utilized the ROSMAP cohort to evaluate RNA sequencing of the dorsal lateral prefrontal cortex (DLPFC) in connection with cognitive ability assessments performed on these individuals prior to death.^58^ A co-expression gene-module associated with cognitive decline was identified with P2RX7 appearing as a high-scoring gene in the module. P2RX7 expression is also found to specifically track with Braak staging, with Braak stage V-VI prefrontal cortex exhibiting significantly increased P2RX7 compared to stages 0-II.^53^ This same study also determined that P2RX7 protein increases coincided with synapse loss in human AD brain samples. These studies demonstrate the key role in AD progression and pathology that places it as a favorable therapeutic target which requires further characterization.

Altogether, previous work has demonstrated P2RX7 has a pivotal role in neuroinflammation and neurodegeneration, and its elevated expression in human AD brain samples points to a critical role in disease pathogenesis. In the present study, our goal is to characterize the immunomodulatory properties of P2RX7 activation in a human microglia-like cell model. Significant differences exist between human and mouse P2RX7 characteristics, emphasizing the requirement for study in human cells for more clinically relevant outcomes. Alternative splicing of P2RX7 in humans yields more splice variants than exist in mice.^59^ Human and mouse P2RX7 also differ in sensitivity to ATP, a foundational function of the receptor, with human P2RX7 displaying higher affinity for ATP.^60^ Here, we add to the growing body of knowledge surrounding P2RX7 function in inflammation and pathogenesis through a human MDMi model reflecting natural human variation in inflammatory function.

## Results

### ATP stimulation in MDMi initiates *IL1β* and *IL18* upregulation

Past studies have shown P2RX7 activation through ATP binding activates the NLRP3 inflammasome and triggers downstream IL-1β and IL-18 production. We cultured monocyte-derived microglia-like cells (MDMi) differentiated from healthy human donors and treated them with 1mM ATP for 3 hours and assessed *IL1β*, *IL18,* and *NLRP3* gene expression. qPCR gene measurement showed increased expression of *IL1β* and *IL18* in MDMi treated with ATP (Figure 1. A). NLRP3 gene expression was not significantly increased by ATP. ATP treatment in MDMi induced *IL1β* and *IL18* gene expression upregulation, which precedes NLRP3 inflammasome activation. When the NLRP3 inflammasome triggers cytokine output, mature cytokine proteins are released from the cell into the extracellular space. We next aimed to measure IL-1β cytokine release from ATP-treated MDMi. Supernatants were isolated from MDMi after ATP stimulation and IL-1β protein was measured using a human IL-1 immunoassay kit (see Methods) that detects proteins using qPCR instrument thermocycling, antibody binding, and fluorescent dyes. Supernatants acquired from ATP-treated MDMi displayed significantly increased levels of IL-1β protein compared to controls (Figure 1. B).

**Figure 1:**
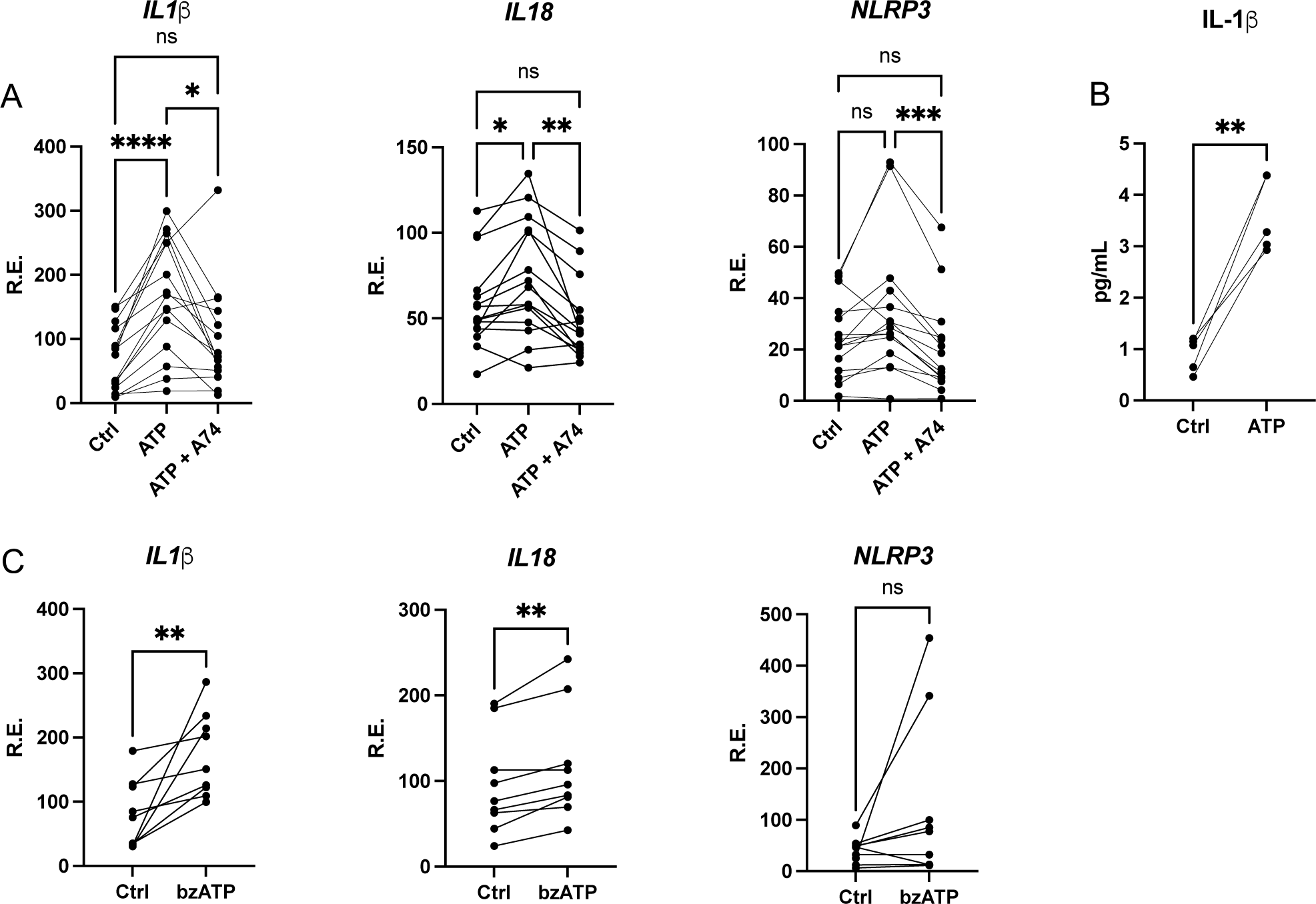
Extracellular ATP Induces *IL1β* and *IL18* gene expression upregulation in a P2RX7-dependent manner. **A)** *IL1β*, *IL18*, and *NLRP3* expression levels were measured with qPCR from MDMi treated with DMSO alone (Ctrl), 1 mM ATP (ATP), or 1 mM ATP and 10 uM A740003 (ATP + A74). Statistics were determined with a repeated measure one-way ANOVA. N= 15. **B)** IL-1β protein detection in control (Ctrl) and ATP-treated (ATP) MDMi supernatants. Statistics were determined with a paired t-test. **C)** *IL1β*, *IL18*, and *NLRP3* expression levels measured with qPCR from MDMi treated with DMSO alone (Ctrl), or 500 uM bzATP (bzATP). Statistics were determined with a paired t-test. N= 10. R.E. (Relative Expression to *GAPDH)*. N=5. ns= p>0.05, *p ≤ 0.05, **≤0.01,***≤0.001, ****≤0.0001

### ATP-driven inflammatory activation is P2RX7-dependent

While elevated eATP levels activate P2RX7, ATP can also bind and activate other purinergic receptors. The selective P2RX7 antagonist A740003 stabilizes the P2RX7 receptor in its closed conformation through allosteric, noncompetitive binding. This binding prevents the downstream effects of eATP.^61–63^ Therefore, the A740003 antagonist was deployed to verify P2RX7-specific cytokine gene expression upregulation. MDMi were preincubated with A740003 for 24 hours followed by three-hour ATP stimulation. Pretreatment with A740003 significantly reduced *IL1β, IL18*, and *NLRP3* gene expression in ATP-treated MDMi (Figure 1. A). Overall, A740003 pretreatment prevented ATP-driven cytokine gene expression changes in MDMi.

We also treated additional MDMi with bzATP, a potent P2RX7-specific agonist to remove the influence of other purinergic signaling in our results. MDMi treated with bzATP exhibited increased *IL1β* and *IL18* expression, further validating this *IL1β* and *IL18* cytokine gene production as P2RX7-specific (Figure 1. C). We conclude *IL1β* and *IL18* gene expression in the presence of elevated eATP levels is P2RX7-dependent in the MDMi model. These experiments demonstrate targeting and activation of the P2RX7 receptor in primary human microglia-like cells by eATP triggers downstream *IL1β* and *IL18* expression and IL-1β protein release.

### ATP reduces MDMi’s ability to uptake Aβ1-42

It is known that P2RX7 in the absence of its ATP ligand and serum can contribute to cell phagocytic abilities.^64^ Whether P2RX7 activation in MDMi mediates the cells’ amyloid-beta uptake capabilities remains to be explored in the human microglia-like cell system. Our results indicate that high-dose ATP reduces Aβ1-42 uptake by MDMi. We found that treatment with ATP reduced the fluorescence signal from HyLite Fluor 647-labeled Aβ1-42 normalized to DAPI. The signal was recovered with the addition of A740003 (Figure 2. A). These results were validated with confocal microscopy where the mean intensity of HyLite Fluor 647-labeled Aβ1-42 was measured in all DAPI-positive cells (Figure 2. B and D, Supplementary Figure 1). Experiments conducted with bzATP treatment also resulted in significant reductions in HyLite Fluor 647-labeled Aβ1-42 uptake, again confirming that activation of P2RX7 with ATP reduces Aβ1-42 uptake (Figure 2. C). These findings align with previous studies demonstrating that the addition of ATP can diminish P2RX7-mediated phagocytic activities.^65^ MDMi treated with A740003 alone showed no change in amyloid-beta uptake (Figure 2. A and Supplementary Figure 2. A). These experiments demonstrate that ATP inhibits Aβ1-42 uptake in human microglia-like cells, and this phenotype is driven by P2RX7.

**Figure 2:**
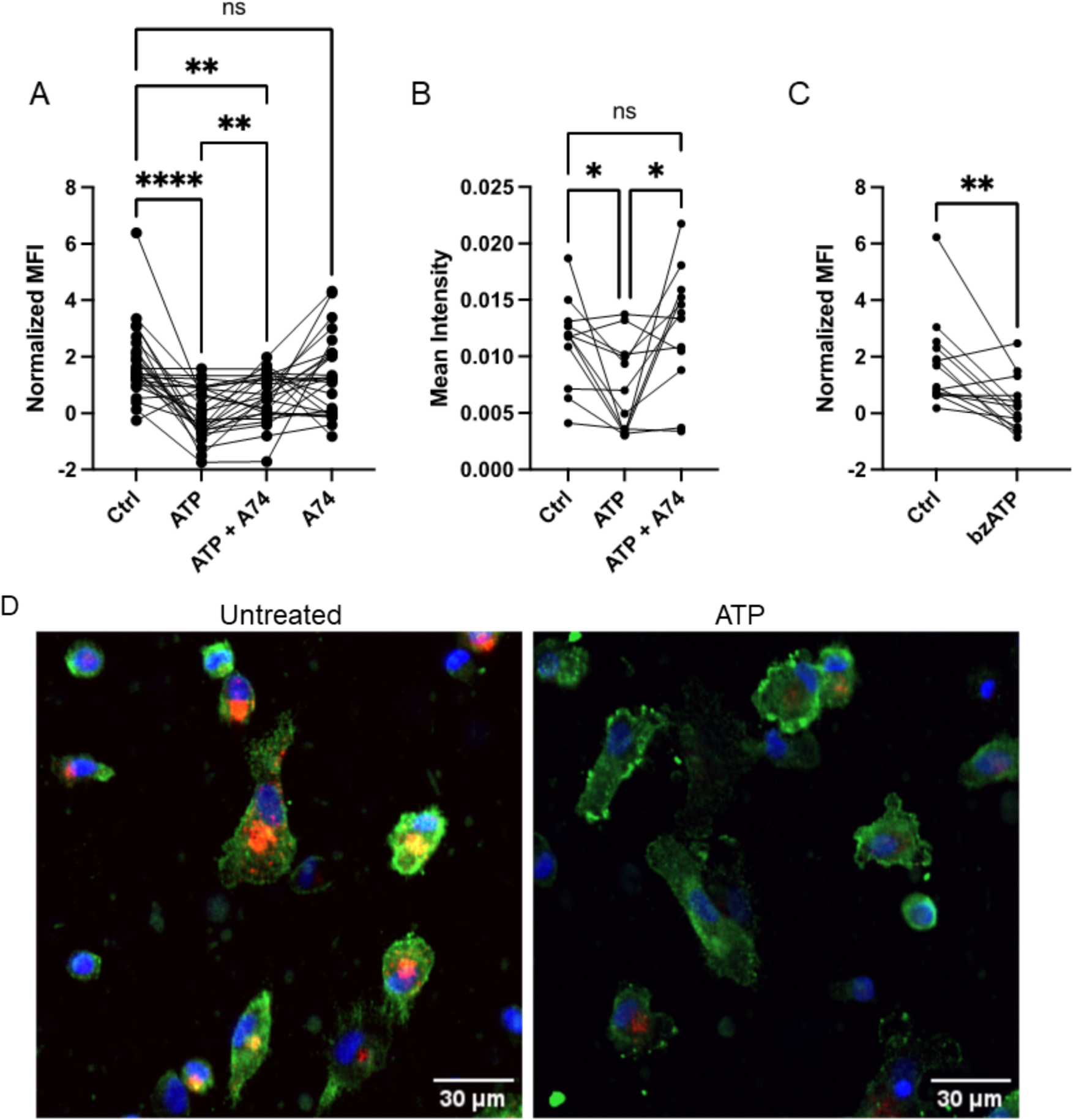
ATP-mediated P2RX7 activation reduces MDMi Aβ1-42 uptake abilities. **A)** Normalized MFI of HyLite Fluor 647-labeled Aβ1-42 peptide uptake normalized to DAPI in MDMi treated with DMSO alone (Ctrl), 1 mM ATP (ATP), 1 mM ATP and 10 uM A740003 (ATP + A74), or 10 uM A740003 (A74). Batch normalization was done in GraphPad PRISM 10 with 0% defined as the smallest mean in each data set and 100% as the average of all means in the data set. Statistics were determined with a repeated measure one-way ANOVA. N=26. **B)** Mean Intensity of HyLite Fluor 647-labeled Aβ1-42 per cell in MDMi imaged using confocal microscopy. MDMi were treated with DMSO alone (Ctrl), 1 mM ATP (ATP), or 1 mM ATP and 10 uM A740003 (ATP + A74). Statistics were determined with a repeated measure one-way ANOVA. N= 12. **C)** HyLite Fluor 647-labeled Aβ1-42 peptide uptake in MDMi treated with DMSO alone (Ctrl), or 500 uM bzATP (bzATP) as measured by fluorescence on a microplate reader. MFI= Mean Fluorescence Intensity normalized to DAPI. Batch normalization was done in GraphPad PRISM 10 with 0% defined as the smallest mean in each data set and 100% as the average of all means in the data set. Statistics were determined with a paired t-test. N=14. **D)** Representative images of MDMi treated with DMSO alone (Ctrl) or 1 mM ATP (ATP) and stained for P2RX7 (green) and DAPI (blue) and used to measure uptake of HyLite Fluor 647-labeled Aβ1-42 peptide (red) and were processed in CellProfiler. ns= p>0.05, *p ≤ 0.05, **≤0.01,***≤0.001, ****≤0.0001

### IL-1RA disrupts ATP activation of P2RX7 in human MDMi

To determine if ATP-mediated reduction in Aβ1-42 uptake is driven by the increase in IL-1β, microglia cultures were pretreated with IL-1RA 24 hours before ATP activation. IL-1RA is a naturally produced antagonist that binds to the IL-1 receptor and acts as a competitive inhibitor for IL-1 binding.^66^ MDMi pretreated with IL-1RA had higher Aβ1-42 uptake levels than ATP alone (Figure 3. A). We next wanted to ask if this IL-1RA-driven uptake increase is accompanied by a reduction in *IL1β* and *IL18* cytokine gene expression. IL-1RA pretreatment significantly reduced *IL1β*, *IL18*, and *NLRP3* gene expression levels compared to ATP alone. Treatment with IL-1RA alone did not significantly change Aβ1-42 uptake (Supplementary Figure 2. B). Altogether, IL-1RA disrupts ATP activation, increases Aβ1-42 uptake levels, and subdues *IL1β*, *IL18*, and *NLRP3* expression in MDMi.

**Figure 3:**
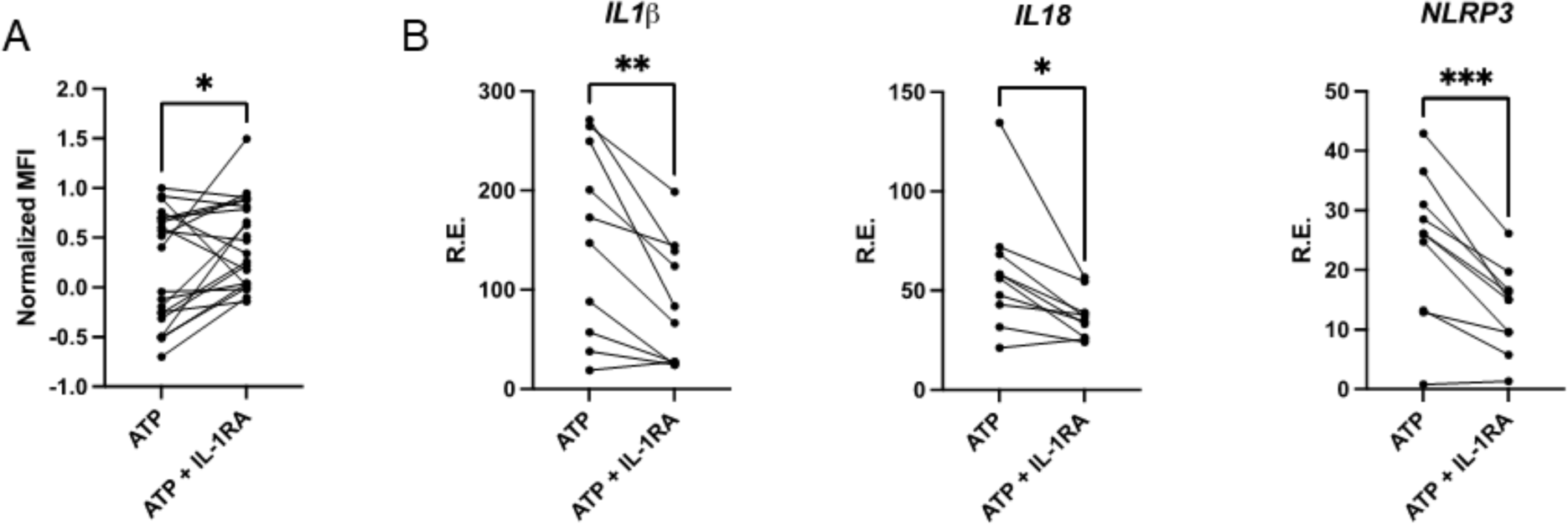
IL-1RA limits ATP modulation of MDMi response. **A)** HyLite Fluor 647-labeled Aβ1-42 peptide uptake in MDMi treated with 1 mM ATP (ATP), or 1 mM ATP and 2.5 ug/mL IL-1RA (ATP + IL-1RA) as measured by fluorescence on a microplate reader. MFI= Mean Fluorescence Intensity normalized to DAPI. Batch normalization was done in GraphPad PRISM 10 with 0% defined as the smallest mean in each data set and 100% as the average of all means in the data set. Statistics were determined with a paired t-test. N=21. **B)** *IL1β*, *IL18*, and *NLRP3* expression levels measured with qPCR from MDMi treated with 1 mM ATP alone (ATP), or 1 mM ATP and 2.5 ug/mL IL-1RA (ATP + IL-1RA). Statistics were determined with a paired t-test. N= 10. R.E. (Relative Expression to *GAPDH)*. ns= p>0.05, *p ≤ 0.05, **≤0.01,***≤0.001, ****≤0.0001

### Influence of donor sex and age on cytokine expression and Aβ1-42 uptake

Aging is the most powerful risk factor for AD and other neurodegenerative diseases.^67^ Sex-dependent differences also play a role in AD risk, with women carrying a higher risk for developing the disease than men.^1,68,69^ These age and sex-driven differences could influence microglia inflammation and neuroimmune mechanisms in disease pathogenesis. Amyloid-beta engulfment and clearance is a central component of clearing the cellular environment and preventing neurotoxicity. We reasoned this process could become impaired in MDMi derived from older donors and may vary between sexes. We therefore aimed to investigate age and sex-driven differences in our MDMi donors’ response to ATP inhibition of Aβ1-42 uptake. There was no significant correlation between Aβ1-42 uptake and age either at baseline or with ATP activation, and no significant difference in Aβ1-42 uptake levels between males and females (Figure 4 A, B). We found no significant differences between female and male donors in *IL1B*, *IL18*, or *NLRP3* gene expression in MDMi at baseline conditions or treated with ATP (Supplementary Figure 3).

**Figure 4:**
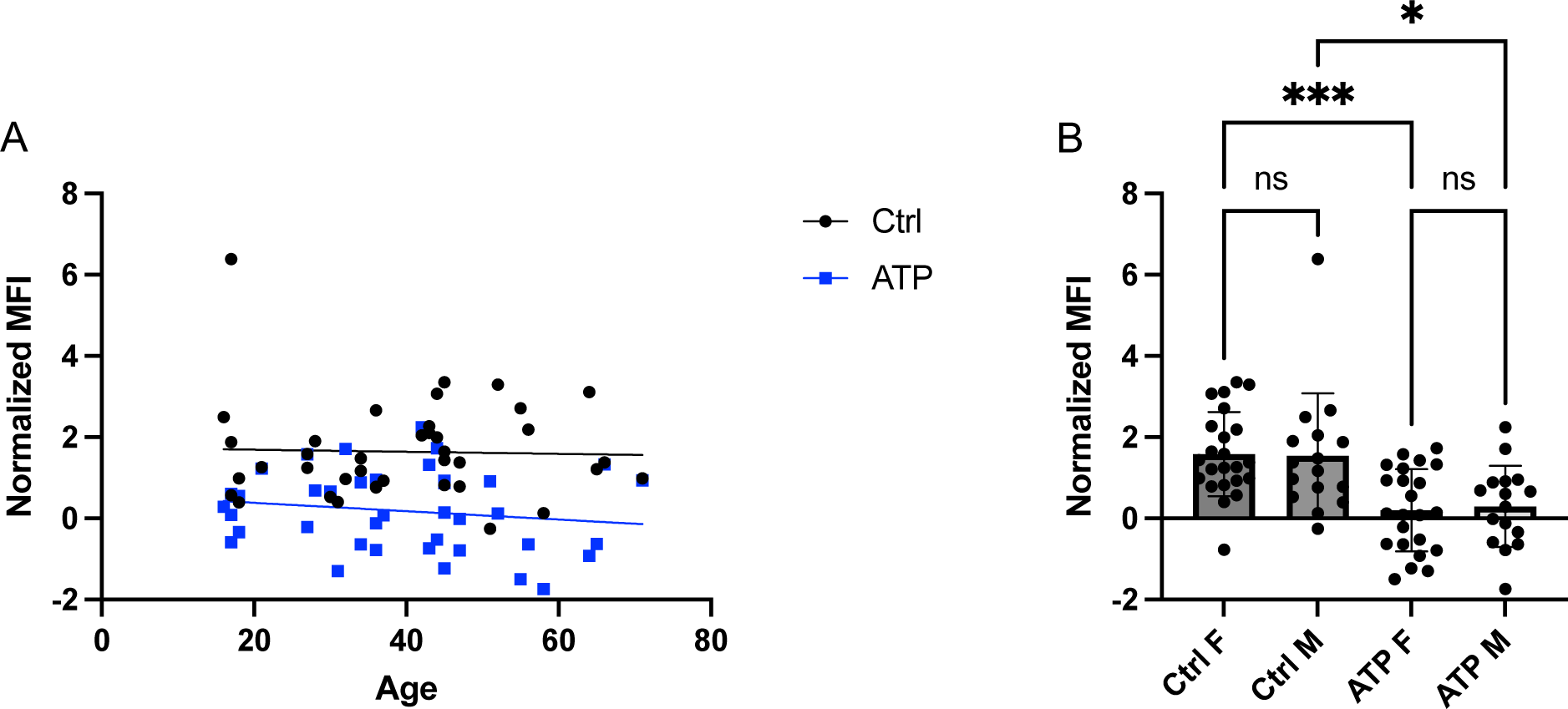
Aβ 1-42 uptake is not modulated by donor age or sex, in control and ATP-stimulated conditions. **A)** Scatter plot of the normalized MFI of HyLite Fluor 647-labeled Aβ1-42 peptide uptake normalized to DAPI in MDMi treated with DMSO alone (Ctrl, black), or 1 mM ATP (ATP, blue) in relation to the age of the MDMi donor. No association with age was determined either in the Ctrl (p=0.85, R^2^=0.001) or ATP (p=0.34, R^2^=0.024) condition. Batch normalization was done in GraphPad PRISM 10 with 0% defined as the smallest mean in each data set and 100% as the average of all means in the data set. Statistics were determined with a simple linear regression. N=39. **B)** Normalized MFI of HyLite Fluor 647-labeled Aβ1-42 peptide uptake normalized to DAPI in MDMi treated with DMSO alone (Ctrl) or 1 mM ATP (ATP) segregated by sex (F=female, M=male). Batch normalization was done in GraphPad PRISM 10 with 0% defined as the smallest mean in each data set and 100% as the average of all means in the data set. Statistics were determined with a mixed-effects analysis. N=23 F and 16 M. ns= p>0.05, *p ≤ 0.05, **≤0.01,***≤0.001, ****≤0.0001

## Discussion

In neurodegenerative diseases, tissue and cellular injury occur, leading to ATP release. The release of ATP into the extracellular environment can occur through dying cells as well as inflammatory signaling.^70–72^ Previous work examining human brain AD neuropathology found elevated P2RX7 expression. Specifically, P2RX7 is upregulated on microglia near amyloid-beta plaques.^54^ Pharmacological inhibition of P2RX7 in rats injected with amyloid-beta 1-42 resulted in neuroprotection.^73^ In a tau mouse model with P2RX7 deleted, spatial memory and synaptic plasticity were rescued.^74^ A deeper understanding of P2RX7’s role in microglia immunomodulation is necessary to interpret how the receptor contributes to AD risk.

In this study, we characterize P2RX7 in a human microglia model derived from multiple human donors, allowing us to investigate potential age and sex differences in ATP-driven P2RX7 activation. We validate ATP-induced *IL1β* and *IL18* upregulation in response to high ATP concentrations and subsequent IL-1β protein release. These pathways are modulated by P2RX7-specific activation as demonstrated through the same pattern shown from MDMi treated with the potent P2RX7 agonist bzATP. Additionally, this cytokine gene expression can be blocked when MDMi are pretreated with the P2RX7 antagonist A740003.

We demonstrated P2RX7 activation with ATP or bzATP in MDMi reduces microglia-like cells’ ability to uptake fluorescent Aβ1-42 in culture. This elimination of phagocytic activity could be averted by blocking ATP activation of P2RX7 with A740003. These experiments confirmed pharmacological inhibition of P2RX7 in human MDMi prevented ATP activation of cytokine expression and loss of Aβ1-42 uptake. We next sought to examine if IL-1 signaling modulated altered phagocytic phenotypes in MDMi by blocking IL-1β’s target receptor, IL-1R. We found pretreatment of MDMi with IL-1RA increased Aβ1-42 uptake levels in MDMi stimulated with ATP. IL-1RA simultaneously reduced *IL1β, IL18*, and *NLRP3* gene expression compared to ATP alone. Taken together, IL-1RA disrupts ATP activation of MDMi and mitigates the proinflammatory phenotype triggered through P2RX7 activation.

P2RX7 is implicated in neurodegenerative diseases beyond AD including MS and ALS. Specifically, elevated P2RX7 levels were found in the activated microglia residing in ALS and MS spinal cords.^45^ In MS patients, two gain-of-function single nucleotide polymorphisms (SNP) in P2RX7 are associated with more severe disease.^47^ In contrast, another P2RX7 variant that leads to loss of receptor function confers a protective effect on risk for MS.^48^ Taken together, there is diverse evidence heavily implicating P2RX7 at the functional and genetic level in a range of neurodegenerative disorders. P2RX7 activity is enhanced in numerous neurodegenerative conditions, making antagonism of the receptor an attractive path toward therapeutics. Blocking P2RX7 activity using antagonists in animal models of PD has been shown to have protective effects.^75,76^ Ongoing studies are characterizing P2RX7 inhibition as a therapeutic target in neurodegeneration.

In the current study, we highlight the ability of P2RX7 inhibition in high ATP conditions in subduing cytokine release and regulating phagocytosis in a human microglia model, suggesting modulation of the receptor in neurodegeneration can point microglia toward a more neuroprotective phenotype.

IL-1RA gained interest as a potential treatment for autoimmune conditions and found success in alleviating symptoms of rheumatoid arthritis in clinical trials.^77^ More recently, Anakinra (drug name of recombinant IL-1RA) was shown to improve hematopoietic regeneration and delay blood aging in mice, solidifying the powerful connection between the immune system and diseases of aging.^78^ The convergence of these pathways places P2RX7-mediated IL-1β release at the crossroads of the immune response, aging, and neurodegeneration.

Whether or not eATP is increased specifically in the vicinity of amyloid beta plaques requires further exploration. Elevated extracellular ATP levels in the brain due to neuronal cell death in AD could inhibit microglia’s ability to clear amyloid beta plaques. As the role of the immune system in neurodegeneration becomes more appreciated, so does the pivotal implications of eATP in the cellular environment. Upregulation of P2RX7 in microglia surrounding plaques suggests this receptor could be sculpting microglial phagocytic and inflammatory responses to aberrant plaques in AD.

Because our experimental model uses human donor blood, there is variation in donor age and sex to consider which may explain why some donors behave differently than others in reaction to ATP. These are important biological factors in the experimental setting as AD is primarily a disease of aging and there are sex-dependent differences in disease risk.^68^ Indeed, there is variability in levels of gene expression and phagocytic responses to ATP in our MDMi cohort, however, the specific trait causing this difference is not clear. We found no association of age or sex with our outcomes. However, it does highlight the clinical relevance of how variability across demographics contributes to disease and the importance of utilizing *in vitro* models to select responders to specific treatments. While no associations were found in our experiments, this does not rule out P2RX7 or ATP-specific differences in age and sex in other disorders. Sex-specific differences have been explored in P2RX7 variation and bipolar disorder, with associations found in females but not males.^79^

Overall, this study provides further characterization of ATP activation and inflammatory pathway modulation by P2RX7. We find ATP’s downstream effects are partially driven by IL-1 signaling and blocking IL-1 signaling through IL-1RA diminishes this proinflammatory phenotype and reestablishes a phagocytic microglia response. P2RX7 and its downstream pathways are significant targets for therapeutic development in AD. Specifically, fine-tuning microglia responses through these pathways may allow for increased pathogenic amyloid-beta clearance and neuroprotection in the aging brain.

## Abbreviations List

ATP: Adenosine triphosphate
eATP: Extracellular ATP
bzATP: 2ʹ(3ʹ)-O-(4-Benzoylbenzoyl)adenosine-5ʹ-triphosphate tri(triethylammonium) salt
P2RX7: Purinergic Receptor 7
IL-1β: Interleukin 1β
IL-18: Interleukin 18
NLRP3: NLR family pyrin domain containing 3
IL-1Ra: Interleukin-1 receptor antagonist

## Methods

MDMi are created as described previously.^80^ Blood from healthy human donors is separated using the density gradient medium Lymphoprep (Stemcell technologies #07851) to isolate mononuclear cells. These peripheral blood mononuclear cells (PBMCs) are cryopreserved in Fetal Bovine Serum (FBS) with 10% Dimethyl sulfoxide (DMSO) at −80°C. Each donor age and sex are recorded if available. PBMCs are thawed and purified monocytes are obtained through CD14+ microbead isolation (Miltenyi #130-050-201). Monocytes are then plated in 96-well plates at a density of 150,000 cells per well and cultured in serum-free RPMI (Gibco #R8758) media with 1% penicillin and streptomycin (10,000 U/mL) (Fisher Scientific 15-140-122) and 2.5 ug/mL Fungizone (Cytiva #SV30078.01). A cytokine mix is added to the media and consists of macrophage colony-stimulating factor (M-CSF) (10 ng/mL), granulocyte-macrophage colony-stimulating factor (GM-CSF) (10 ng/mL), nerve-growth factor-β (NGF-β) (10 ng/mL), chemokine ligand 2 (CCL2) (100 ng/mL), and interleukin-34 (IL-34) (100 ng/mL). Cytokines were purchased from R&D Systems (NGF-β, GM-CSF, and IL-34) and Biolegend (M-CSF and CCL2). These cytokines facilitate microglia development and differentiation into monocyte-derived microglia-like cells (MDMi).^81–83^ Monocyte-derived microglia-like cell models have been reviewed and characterized as an appropriate model to study human microglia *in vitro*.^84^

### ATP and bzATP stimulation

MDMi on day 11 of differentiation are treated with 1 mM ATP or 500 uM bzATP for 3 hours at 37°C. As both are resuspended in DMSO, DMSO is added to controls. ATP was purchased from Theromfisher (#R0441) and bzATP was purchased from Tocris Bioscience (#3312).

### A740003 and IL-1RA treatment

The A740003 P2RX7 antagonist (Tocris biotechne, #861393-28-4, Batch #3B/266155) is dissolved in DMSO and added to MDMi cultures at a concentration of 10 uM. The recombinant human IL-1RA protein was purchased from R&D Systems (#280-RA-050/CF) and reconstituted in phosphate-buffered saline (PBS). IL-1RA was added to cultures at a concentration of 0.5 ug per well (2.5 ug/mL).

### Cell Lysis and RNA isolation

MDMi are lysed with RLT buffer (Qiagen #74104) with 1:100 β-mercaptoethanol, purified RNA is extracted using the RNeasy 96 well plate isolation kit (Qiagen #74182).

### Reverse Transcriptase Polymerase Chain Reaction

RNA is transformed to cDNA using reverse transcription PCR. The PCR mix consists of dNTP mix (#R72501), Random Hexamers (#N8080127), RNase Inhibitor (#N8080119), MgCl_2_ (#AB0359), 10x PCR buffer (#4486220), and M-MLV Reverse Transcriptase (#28025013) purchased from Thermofisher Scientific. Reagents and RNA are loaded into a 96-well plate to a total volume of 50 uL and run in an Applied Biosystems MiniAmp Thermocycler. Thermocycler program: 25°C for 10 minutes, 48°C for 45 minutes, 95°C for 5 minutes, and held at 4°C upon run completion.

### Quantitative Polymerase Chain Reaction and Primers

Sample cDNA is loaded into a 96-well PCR plate with Taqman Fast Advanced Mastermix (#4444554), assay primer for the target gene to be detected with FAM, and housekeeping gene to be detected with VIC. qPCR primers were purchased from Thermofisher Scientific (GAPDH: Hs02786624_g1, IL1B: Hs01555410_m1, IL18: Hs01038788_m1, NLRP3: Hs00918082_m1, P2RX7: Hs0017521_m1). For all experiments, the housekeeping gene glyceraldehyde-3-phosphate dehydrogenase (GAPDH) was used to normalize all values to relative expression (R.E.). Samples are run with the protocol: Stage 1 (x1): 50°C for 2 minutes, 95°C for 2 seconds. Stage 2 (40x): 95°C for 1 second, 60°C for 20 seconds. Cycle threshold (Ct) values are collected and normalized to GAPDH Ct.

### Protein Immunoassay

IL-1β protein levels were quantified using Thermofisher Scientific Invitrogen ProQuantum Human IL-1 Immunoassay Kit (#A35574). MDMi supernatants were removed from culture plates and cryopreserved at −80°C until use for assay. Briefly, Antibody-conjugate A and Antibody-conjugate B were combined with conjugate dilution buffer. Each supernatant sample was diluted 1:1 in Assay Dilution Buffer. Serial dilutions of the reconstituted standard were made for the creation of a standard curve ranging from 0 pg/mL to 1,000 pg/mL. To bind the analyte, diluted samples or standards were combined with the antibody-conjugate mixture in a 96-well plate and incubated for 1 hour at room temperature. After this incubation, Master Mix and Ligase were added to all reaction wells. The PCR protocol was run on an Applied Biosystems instrument with the following protocol: Ligation: 25°C for 20 minutes; Ligase inactivation: 95°C for 2 minutes; Denaturation and Annealing/extension (40 cycles): 95°C for 15 seconds followed by 60°C for 1 minute (40 cycles). The data was analyzed using Invitrogen Proquantum cloud-based software.

### Aβ 1-42 uptake

MDMi treated with various conditions were incubated with 10 uM HyLite Fluor 647-labeled Aβ1-42 peptide (AnaSpec) for 1 hour at 37°C.

### Immunocytochemistry

Media is removed from wells and cells are washed with PBS. Cells are then fixed with 4% Paraformaldehyde (PFA) for 15 minutes on ice. PFA is removed and cells are PBS washed before staining with Anti-P2X7 Receptor (extracellular)-FITC antibody purchased from Alomone Labs (#APR-004). This conjugated antibody is incubated on the cells overnight before washing and addition of DAPI NucBlue Fixed Cell Stain ReadyProbes (#R37606) the following day.

### Fluorescence measurements on plate reader

Fixed cell fluorescence was measured in black 96-well plates using a Tecan Infinite F200 pro fluorescent plate reader. Mean Fluorescence intensity (MFI) was measured and HyLite Fluor 647-labeled Aβ1-42 fluorescence was normalized to total DAPI fluorescence to normalize for the number of cells per well.

### Microscopy

Cells were imaged on the Andor BC43 Oxford Instruments benchtop microscope using a programmed high throughput imaging method. Images were processed using FIJI image processing software and analyzed using the open-source program CellProfiler to classify and count cells.

### Statistics

Statistics were performed using Prism (GraphPad Software). Student’s t test, one-way ANOVA, simple linear regression, or mixed-effects analysis were calculated. Normalization was also done in Prism for batch correction. Statistical significance was classified as P values less than 0.05. ns= p>0.05, *p ≤ 0.05, **≤0.01,***≤0.001, ****≤0.0001.

## Acknowledgements

The authors are grateful to the participants of the NYBC for the time and specimens that they contributed. We would like to thank Carol Troy for the use of her microscope. This work was supported by the US National Institutes of Health grants RF1AG058852-01, RF1AG058852-01S1, R01AG076018-01 and R21AG073882-01. The content is solely the responsibility of the authors and does not necessarily represent the official views of the National Institutes of Health.

## Author Contributions

K.S.H and EMB designed the experiments and wrote the manuscript. All authors read and edited the manuscript. K.S.H. conducted the gene expression and phagocytosis experiments. K.S.H. and K.K. conducted the IL-1β protein immunoassay experiments. M.Y. organized and recorded NYBC demographic information.

