## Supplementary Data for "IL-1RA Disrupts ATP Activation of P2RX7 in Human Monocyte-Derived Microglia-like Cells"

### Manuscript Supplementary Figures:

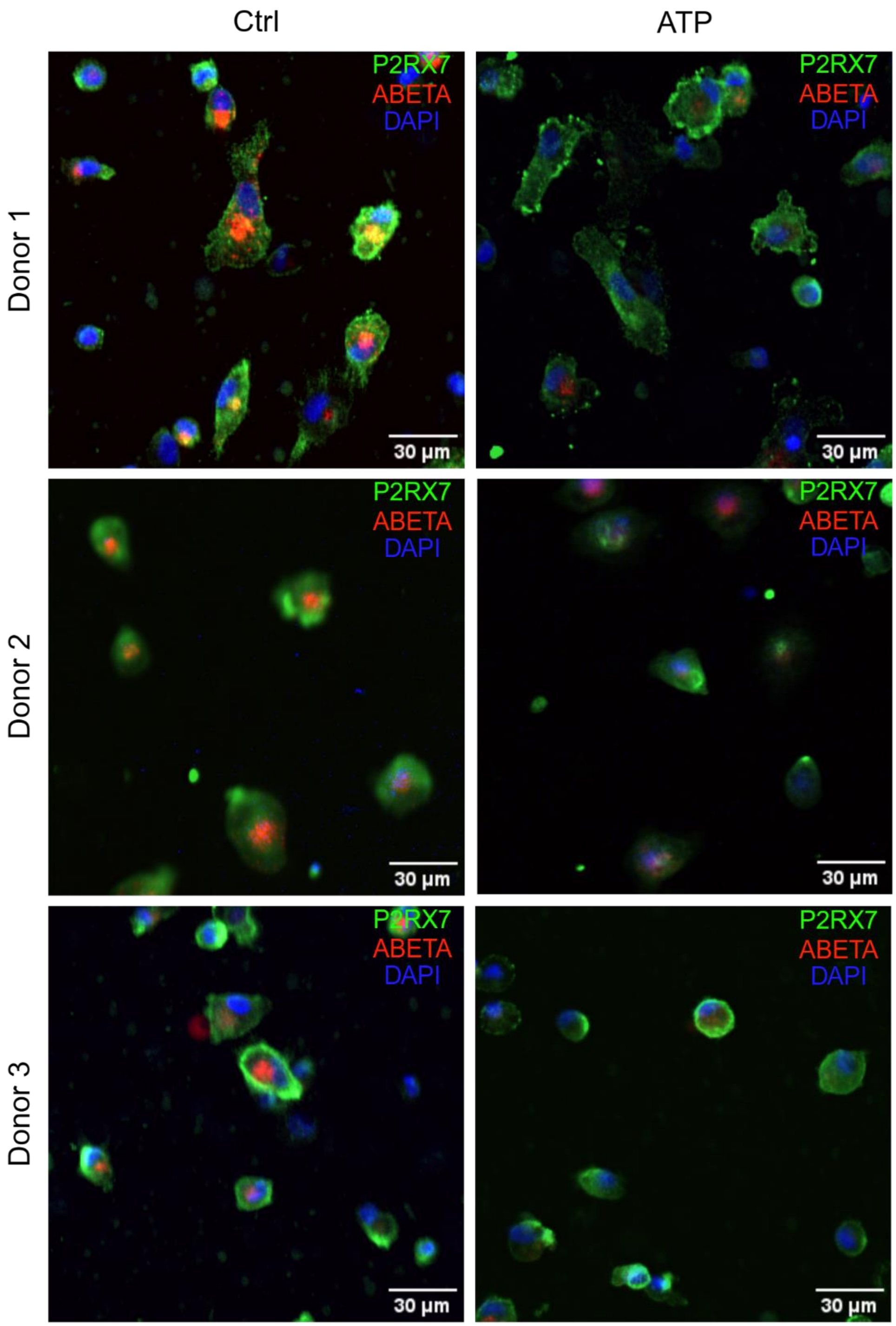


**Supplementary Figure 1: Representative images of MDMi donors in untreated and ATP-stimulated cultures.** Representative images of MDMi treated with DMSO alone (Ctrl) or 1 mM ATP (ATP) from three different MDMi donors and stained for P2RX7 (green) and DAPI (blue), and used to measure uptake of HyLite Fluor 647-labeled Aβ1-42 peptide (red). Images were processed in CellProfiler.

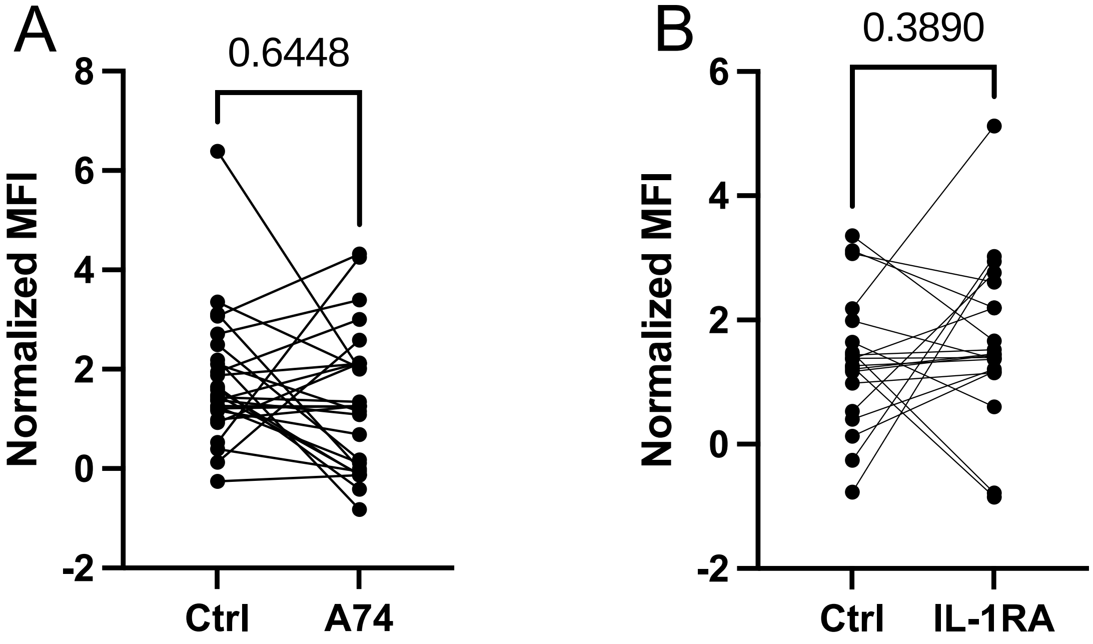


**Supplementary Figure 2: The addition of A740003 or IL-1RA to MDMi cultures does not modulate Aβ1-42 uptake. A)** Normalized MFI of HyLite Fluor 647-labeled Aβ1-42 peptide uptake normalized to DAPI in MDMi treated with DMSO alone (Ctrl) or 10 uM A740003 (A74). Batch normalization was done in GraphPad PRISM 10 with 0% defined as the smallest mean in each data set and 100% as the average of all means in the data set. Statistics were determined with a paired t-test. N=26. **B)** Normalized MFI of HyLite Fluor 647-labeled Aβ1-42 peptide uptake normalized to DAPI in MDMi treated with DMSO alone (Ctrl) or 1 mM ATP and 2.5 ug/mL IL-1RA (ATP + IL-1RA). Batch normalization was done in GraphPad PRISM 10 with 0% defined as the smallest mean in each data set and 100% as the average of all means in the data set. Statistics were determined with a paired t-test. N=20. ns= p>0.05

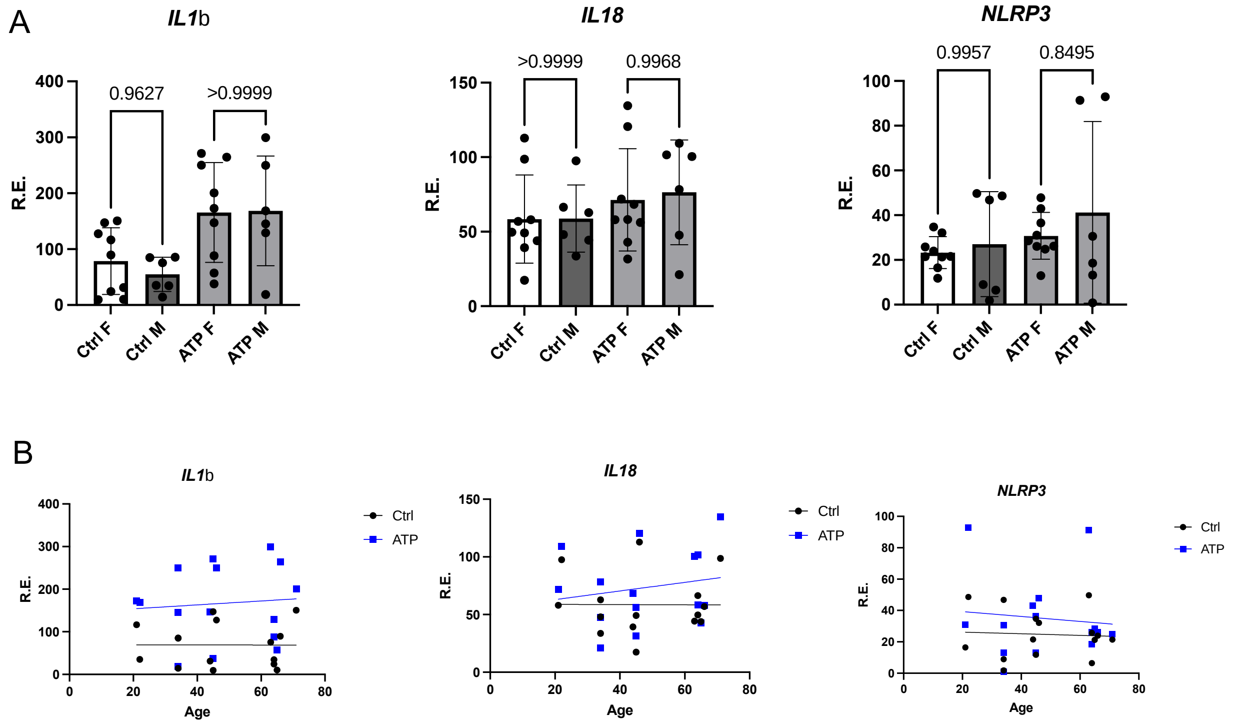


**Supplementary Figure 3: *IL1β*, *IL18*, and *NLRP3* expression is not modulated by donor age or sex, in control and ATP-stimulated conditions. A)** *IL1β*, *IL18*, and *NLRP3* expression levels were measured with qPCR in MDMi treated with DMSO alone (Ctrl) or 1 mM ATP (ATP) segregated by sex (F=female, M=male). There was no significant difference between males and females in either control or ATP-treated MDMi. Statistics were determined with a mixed-effects analysis. N=9 F and 6 M. ns= p>0.05 **B**) Scatter plot of *IL1β*, *IL18*, and *NLRP3* expression levels in MDMi treated with DMSO alone (Ctrl, black), or 1 mM ATP (ATP, blue) in relation to the age of the MDMi donor. No association with age was determined either in the Ctrl or ATP condition. *IL1β*: Ctrl (p=0.99, $R^2$=8.7x10^-6) or ATP (p=0.76, $R^2$=0.007); *IL18:* Ctrl (p=0.99, $R^2$=2.3x10^-5) or ATP (p=0.51, $R^2$=0.034); *NLRP3:* Ctrl (p=0.84, $R^2$=0.003) or ATP (p=0.72, $R^2$=0.01). N=15.
